# Estimation of free-roaming dog populations using Google Street View: A Validation Study

**DOI:** 10.1101/2024.06.03.596211

**Authors:** Guillermo Porras, Elvis W. Diaz, Micaela De la Puente, Cesar M. Gavidia, Ricardo Castillo-Neyra

## Abstract

Free-roaming dogs play a central role in carrying zoonotic pathogens such as rabies virus, *Echinococcus granulosu*s, and Leishmania spp. The control and elimination of these pathogens require quantitative knowledge of dog populations. Thus, estimating the dog population is fundamental for planning, implementing, and evaluating public health programs. However, dog population estimation is time-consuming, requires many field personnel, may be inaccurate and unreliable, and is not without danger. Our objective was to validate a remote methodology for estimating the population of free-roaming dogs using Google Street View (GSV). Our target populations were free-roaming dogs from Arequipa, Peru, a rabies-affected area. Adopting a citizen science approach, and using social media, we recruited online citizen scientists from Arequipa and other regions and trained them to use GSV to identify and count free-roaming dogs in 26 urban and periurban communities. We used correlation metrics and negative binomial models to compare the counts of dogs identified in the GSV imagery with accurate counts of free-roaming owned dogs estimated via door-to-door surveys. In total, citizen scientists detected 868 dogs using GSV and using door-to-door surveys we estimated 909 free-roaming dogs across those 26 communities (Pearson’s coefficient was r=0.73, p < 0.001). Our model predicted that for each free-roaming dog detected with GSV in urban areas, there were 1.03 owned dogs with free access to the street (p < 0.001). The type of community, urban versus periurban, did not have an important effect on the model, but fitting the models in periurban communities was difficult because of the sparsity of high-resolution GSV images in those areas. Using GSV imagery for estimating dog populations is a promising tool in urban areas. Citizen scientists can help to generate information for disease control programs in places with insufficient resources.

## Introduction

Free-roaming dogs are a public health problem because of the transmission of diseases that affect other animals (1–6) or people (7). Developing countries are the most affected areas with dog-transmitted diseases (7,8). The global dog population has been estimated at more than 990 million, according to the Wildlife Conservation Society (WCS), of which approximately 700 million are free-roaming dogs (9). The free-roaming dog population is composed of owned dogs that are allowed outside houses without restriction, previously owned dogs that escaped, were abandoned, or got lost, and dogs born ownerless. Detailed information on these dog subpopulations is crucial for programs aimed at eliminating dog-mediated diseases. Planning, implementation, and evaluation of control strategies are hampered by the lack of information about free-roaming dog population size (10).

Methods recommended by the World Health Organization (WHO) to estimate free-roaming dog populations include 1) total or direct counts; 2) estimation method by catch rates; 3) method of recapture; and 4) the Beck method or photographic recapture method (11). Other methods have also been reported: 5) Mark-resight procedures, 6) a simple count of the observed dogs and the rate of encounters (individuals/km2), 7) dogs marked at the time of vaccination followed by home visits and monitoring of the animal tag in the target area, 8) dogs marked at the time of vaccination followed by home visits and monitoring of the animal tag in the target area, and 9) Pasteur technique (12). These methods have important drawbacks, such as high variability of results (12,13), being time-consuming like in the method of direct counts (13), and requiring trained personnel (14,15). In addition, the implementation of methods like surveys can have high costs (16). Program costs are particularly concerning in resource-constrained settings, and wherever inadequate resources are assigned to public health strategies responsible for managing and preventing the spread of dog-mediated diseases. There is a need to develop new methods to estimate free-roaming dog populations.

One potential alternative is to use an already existing and easily accessible database like the images captured by Google Street View (GSV). GSV is made up of billions of images to display a virtual representation of our world on Google Maps and Google Earth, allowing users to explore the world virtually and its content comes from Google and its partners (17). This technology provides virtual panoramas using an advanced camera system mounted on moving vehicles; these vehicles revisit areas to update the imagery (18). The available up-to-date imagery and the ease of use of GSV may facilitate its application in digital epidemiology studies (19–21). GSV imagery has been successfully used to verify structures in field audits of neighborhoods (22), train artificial intelligence by counting pedestrians (23), identify patterns of traveling to understand, plan and evaluate policies and interventions in urban mobility (24), analyze the distribution of moths in trees for ecological research (25), assess the nesting habitat of vultures for conservation purposes (26), and evaluate the association between the characteristics of the neighborhoods and the cases of COVID-19 (27). However, its potential for free-roaming dog population estimation has not yet been fully explored.

Our aim was to evaluate a remote methodology for estimating the free-roaming canine population using GSV imagery as compared with previous data gathered by door-to-door surveys. For rabies, knowledge of the free-roaming dog population is crucial since these dogs have higher contact rates and may be responsible for most of the transmission and persistence of the rabies virus (28). Efforts to eliminate canine rabies should pay special attention to this subpopulation of dogs (29). Developing a methodology to estimate the free-roaming dog population could be useful in countries where the dog population size is unknown and other methods are unfeasible (15).

## Methods

### Study Area

We conducted this study in Arequipa City, southern Peru, where canine rabies has re-emerged as a worrying threat since 2015 (30). Specifically, this study was conducted in the district of Alto Selva Alegre (ASA) (human population for 2016: 83,310). ASA has had canine rabies cases since 2015 (31,32). We used GSV images captured in the province of Arequipa, located in the South of Peru and situated at ∼2,300 meters above sea level, with a population of 969,000 people. A total of 26 communities (administrative divisions within a district) were included in the study, 20 in urban areas and 6 in periurban areas (Fig 1). The availability of GSV images is higher in urban areas of the city, due to the greater number of paved streets that allow the accessibility of the cars that capture the images.

**Fig 1.**
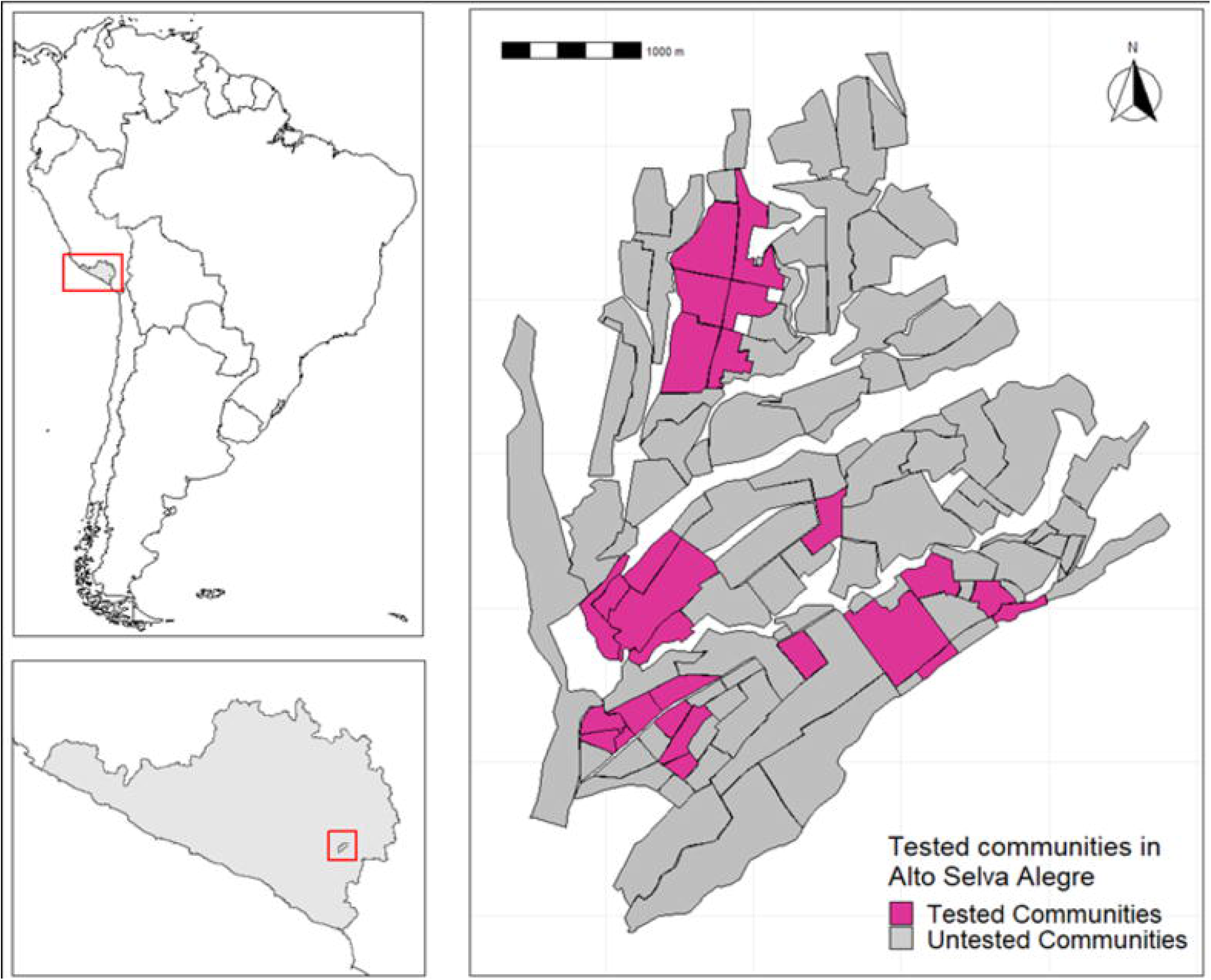
Study communities in the district of Alto Selva Alegre, Arequipa, Peru.

### Door-to-door household surveys

In 2016, the Laboratory of Research in Zoonotic Diseases of the Universidad Peruana Cayetano Heredia conducted door-to-door surveys in the district of ASA, in 42 communities (21 urban and 21 periurban). Each household was visited, and information was collected on the house characteristics, the dog owner or interviewee, and the dog, aiming to obtain demographic data on the dog population. All houses in the study communities were geocoded and survey data were linked to household coordinates. These surveys helped to estimate the total owned dog population and the owned free-roaming dog population in every community.

### Citizen scientists recruitment and training on GSV imagery search

Twenty-two citizen scientists were recruited via social networks through publications on Facebook pages related to veterinary medicine, veterinary schools, and animal protection. In addition, we invited people from the School of Veterinary Medicine at Universidad Nacional Mayor de San Marcos through WhatsApp messages. All citizen scientists received the following materials: i) a digital manual with the search protocol; ii) an Excel file for registering the data of the dogs detected; iii) the link of the assigned area (polygon drawn on Google Maps); iv) and the date of the images captured in GSV. All citizen scientists received the search protocol via email with clear instructions to implement the search in GSV and collect the data. Whatsapp sessions were used to solve doubts and respond to questions from citizen scientists during the COVID-19 pandemic restrictions. The training for the search and counting of free-roaming dogs using GSV was performed taking as a basis the World Society for the Protection of Animals (WSPA) protocol (33). Citizen scientists were instructed to search using the images dated September 2013, which comprised most of the available images. Also, they received the indication to perform the search using GSV covering all streets shown on the map where GSV imagery is available within the assigned community. The collected information was submitted by email and corroborated by a proficient researcher.

### Piloting GSV imagery search

We conducted a pilot study to first, develop the appropriate protocol for free-roaming dog search in designated areas and assess the detection capability of citizen scientists to follow. Seven members of our team and 28 citizen scientists searched free-roaming dogs in 73 urban and periurban communities of Arequipa City (different than the 26 communities used in the final analysis of this study). The protocol was refined during this process and it was clear that, on average, one citizen scientist would miss a substantial number of free-roaming dogs and multiple citizen scientists per area would improve detectability and accuracy. Similar to other studies that employed a citizen science approach to detect events over space (26), our next step was to determine the optimal number of citizen scientists per area/community.

### Determining the appropriate number of citizen scientists per community

We conducted a preliminary study to determine the optimal number of citizen scientists needed within a community to conduct searches that produced an accurate number of free-roaming dogs. Out of the 73 communities searched during the pilot, 6 were chosen randomly. In those 6 communities, 22 citizen scientists were randomly assigned to a community, until each community had 4 searchers. The 4 searchers in each community, look for dogs over the same areas and streets, but they were blinded from their peers’ results. The six communities were also searched by the proficient investigator who served as a goal standard. We compared the number of unique dogs detected by 1, 2, 3, and 4 citizen scientists in the same community versus the number of dogs detected by the proficient investigator.

### Dog detection in GSV imagery

After the optimal number of searchers per community (4) was determined, the 22 citizen scientists were randomly assigned to one of the 26 final study communities, allowing for repetition until each community had 4 citizen scientists to search for dogs. All the sets of 4 searchers per community were different. The 4 citizen scientists in each community, look for dogs over the same areas and streets, but they were blinded from their peers’ results. The search was performed between June and December of 2020. The citizen scientists scanned images in a virtual walking tour (Fig. 2), and at each stop point, they turned laterally 360°. All forms (shapes) that looked similar to a free-roaming dog and all dogs clearly identified were registered. If the same dog appeared in different images in the same street, the citizen scientists only sent one link image. If the citizen scientists were unsure if the dog found was duplicated, it was considered a different dog and the image links were sent, and registered in the Excel file. Each time a free-roaming dog was identified, citizen scientists completed the following variables in the Excel sheet: i) the month and year when Google captured the image; ii) the community where the search was conducted; and iii) the image link. When more than one dog was identified in the same image, the same registration protocol was conducted for each dog. Only dogs unleashed and walking freely in the street were counted (Fig 3). The citizen scientists emailed the Excel database to the proficient investigator to evaluate the images.

**Fig 2.**
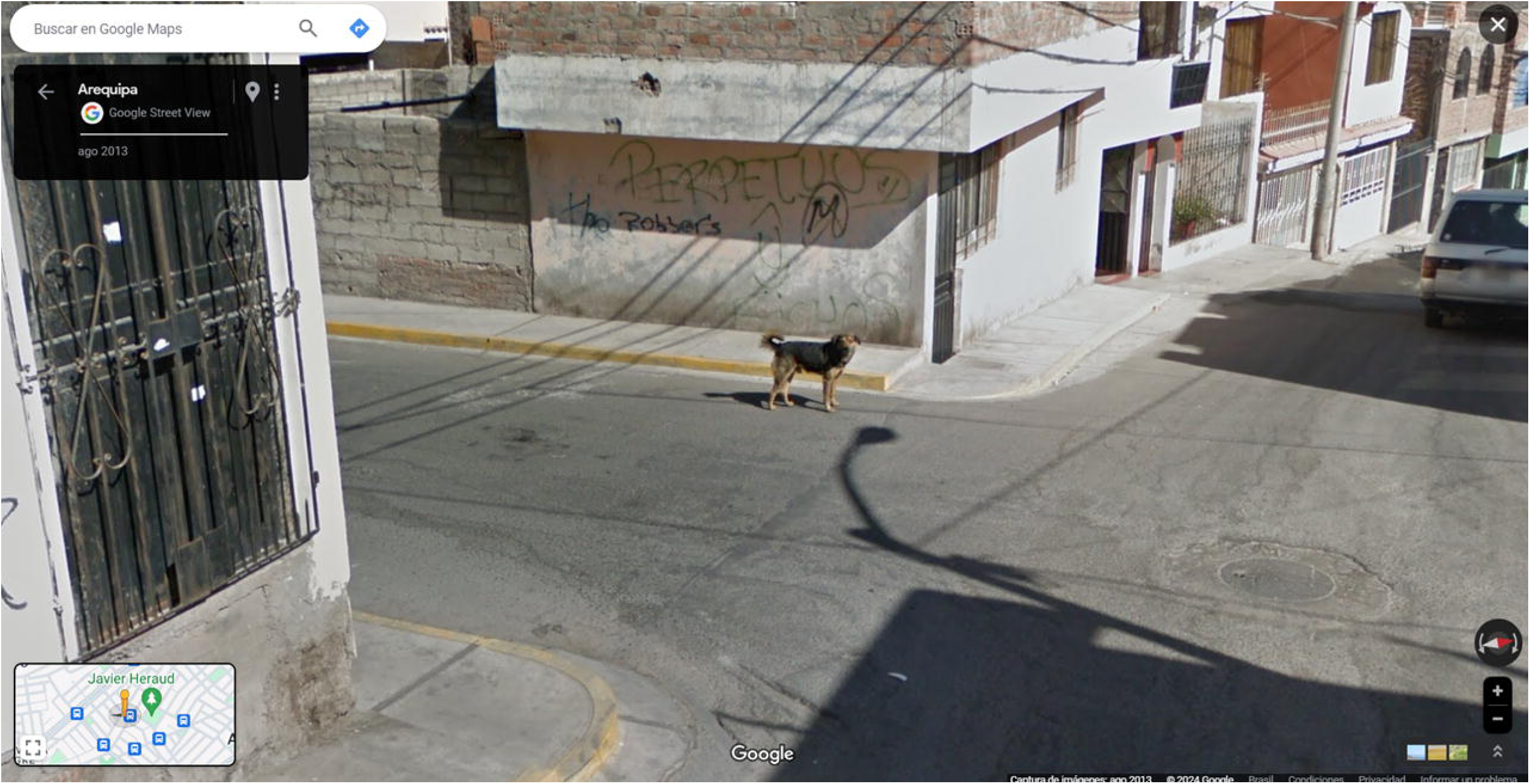
Free-roaming dog detected using GSV imagery. Image obtained from Google Street View (Google Maps™, © Google) (Accessed: 2024-04-27)

**Fig 3.**
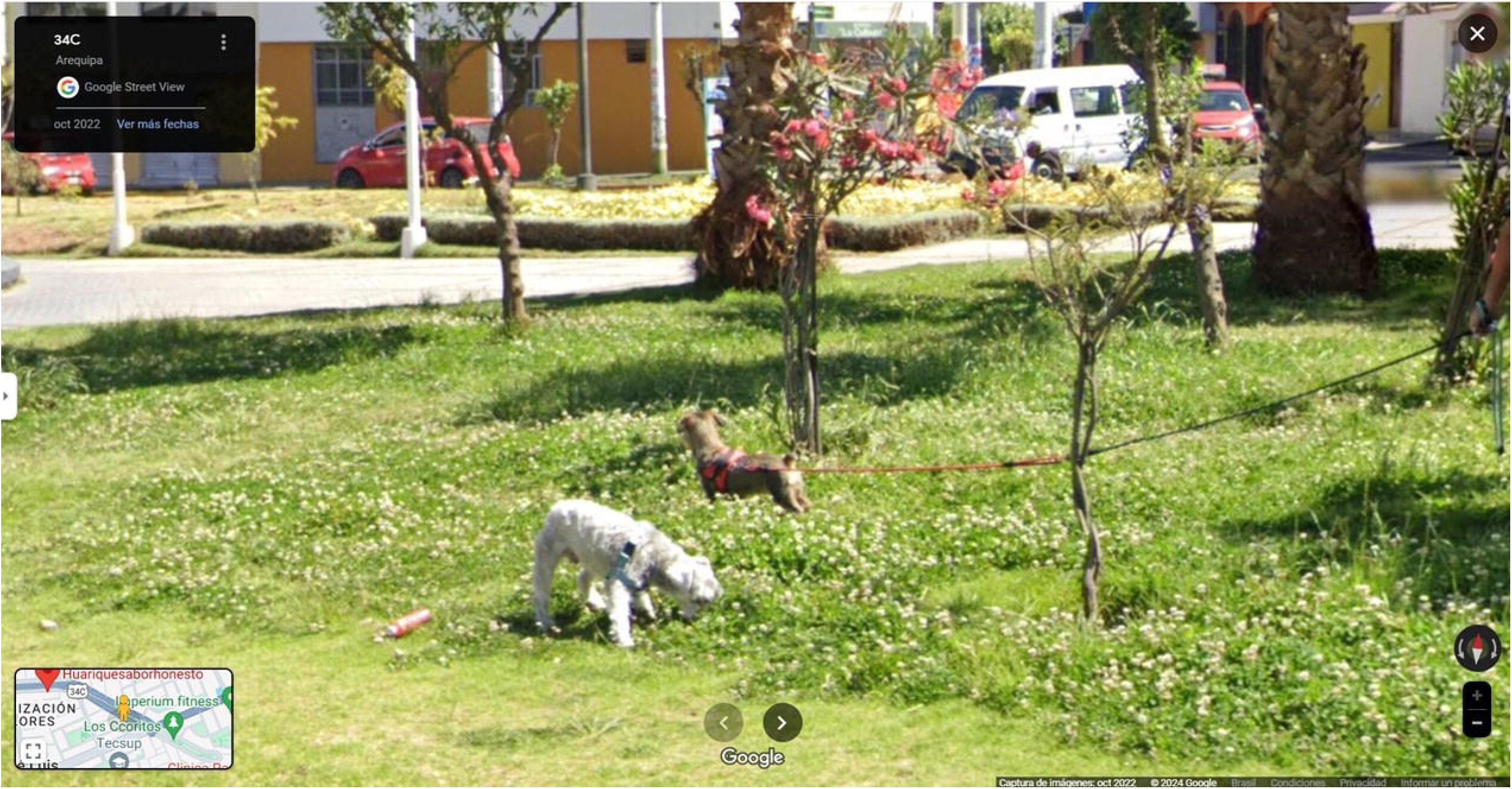
GSV imagery showing dogs on the leash. These dogs were not included in the final database. Image obtained from Google Street View (Google Maps™, © Google) (Accessed: 2024-04-27)

### Data quality control

The data collected by the citizen scientists were validated by a proficient investigator (GP). The verification process involved a meticulous examination of each GSV image through the hyperlinks preserved by citizen scientists in the Excel database. The proficient investigator determined whether the purported detection took place on the specified date, verified the status of detected dogs as free-roaming, confirmed their presence within the boundaries of the assigned search area of the respective citizen scientist, and addressed instances of potential double-counting by the same citizen scientist within the same search area. Each detected free-roaming dog was systematically coded, georeferenced, and analyzed to compare the similarity between the search results reported by every one of the four citizen scientists assigned to each community.

The number of dogs detected for the ideal number of citizen scientists to search for free-roaming dogs (4) was taken as the maximum number of discoverable dogs using GSV in each of the study communities. Additional data extracted from Google Maps and the surveys were i) The distance traveled during the search (measured in kilometers); ii) the area of each community (measured in hectares); iii) the number of houses in each community; and iv) the classification of the community as either urban or periurban.

### Statistical Analysis

We employed Pearson’s coefficient to assess the correlation between the number of free-roaming dogs found using GSV and the count of dogs with free access to the street in the same communities obtained through door-to-door surveys. For estimating confidence bands around our predicted lines (Fig. 4) we used the function geom_smooth with the R package “ggplot”. We explored different model distributions for the number of free-roaming dogs, such as Poisson, negative binomial, and normal. This distribution of the number of dogs with free access to the street reported in the surveys was assessed using the statistical package “Fitdistrplus” within the R programming environment. This package facilitates the fitting of different statistical distributions to the data, enabling the determination of parameters that optimize the fit. Based on the distribution of the response variable, the final regression model to estimate the association between the predictor variables (dogs detected in GSV, traveled distance, area of the community, number of houses in the community, and type of community), and response variable (free-roaming dogs from surveys) was a negative binomial regression model. For model building, we used a forward stepwise regression approach. Multicollinearity was estimated by calculating the variance inflation factor between the predictor variables. A negative binomial regression was used to build the model, and the number of dogs detected by GSV (discrete) and type of zone (categorical: urban/periurban) were used for modeling. For model selection, we used Akaikes information criteria (AIC) and the significance of the variables included in our model. All the analyses and figures were produced in R (34). All tests were evaluated with a statistical significance level of 0.05.

**Fig 4.**
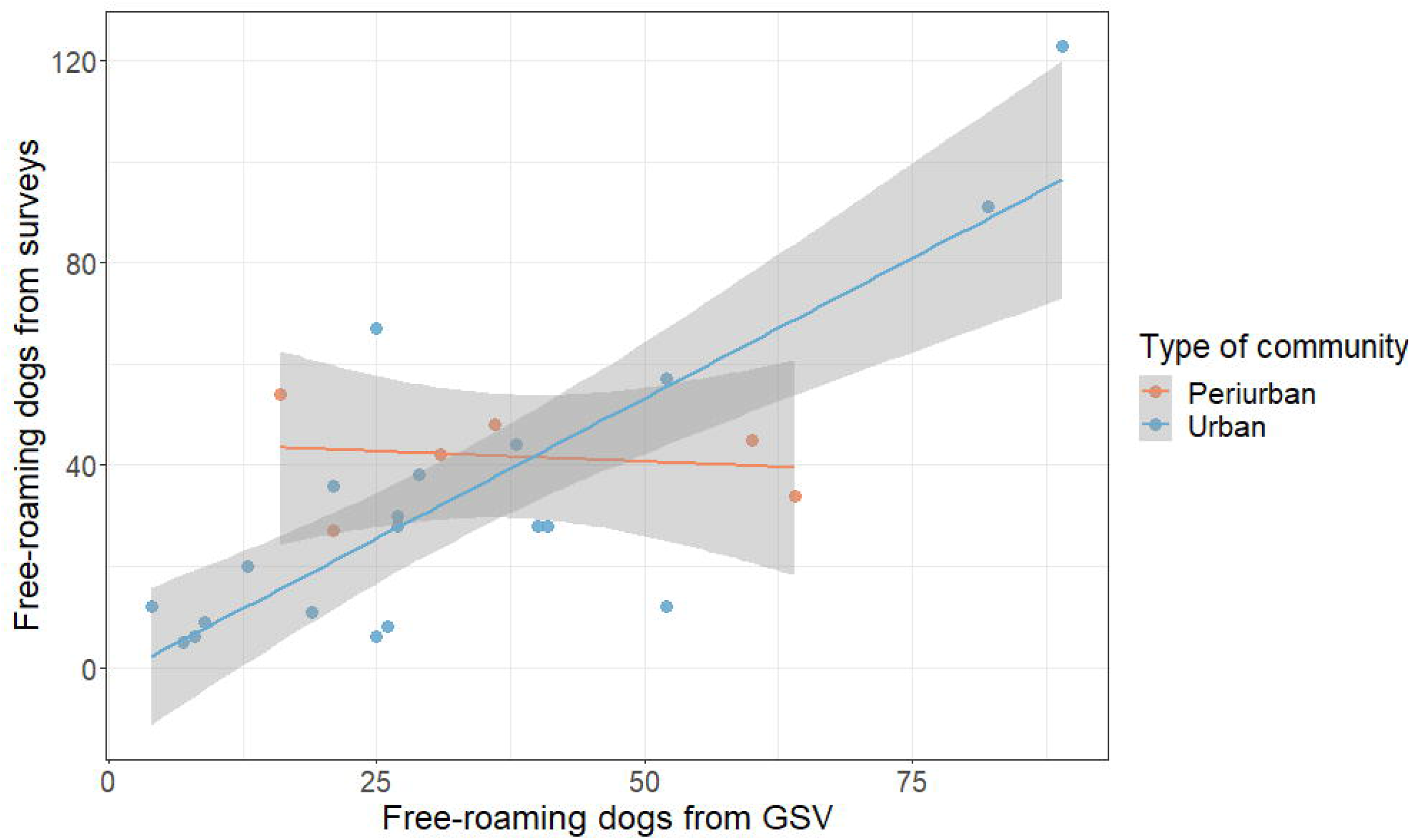
Number of free-roaming dogs detected with GSV per community versus number of free-roaming dogs reported in household surveys in the same communities. Also shown are prediction regression lines and confidence bands.

## Results

### Searching for free-roaming dogs

Using GSV, a total of 868 unique free-roaming dogs were found in 20 urban and 6 periuban communities. The combined area of all communities was 223.3 hectares, and the total length of streets and roads the citizen scientists scanned with GSV was 59.38 kilometers. The 2016 survey registered a total of 909 owned dogs with free access to the street within a total dog population of 4,714. There were marked differences in the number of detected free-roaming dogs and other variables between urban and periurban communities (Table 1). In the community with the highest detection, 89 free-roaming dogs were recorded, while in the community with the lowest detection only 4 were recorded.

**Table 1.** Free-roaming dogs counted with Google Street View and with door-to-door surveys in study community in Arequipa, Peru.

| Variable | Periurban communities (n=6) | Urban communities (n=20) | p-value* |
| --- | --- | --- | --- |
| Number of free-roaming dogs from surveys (median) | 250 (43.5) | 695 (32.95) | 0.136 |
| Number of dogs found in GSV (median) | 228 (33.5) | 634 (31.7) | 0.429 |
| Total length of streets in kilometers (median) | 13.35 (1.97) | 46.03 (2.3) | 0.605 |
| Total study area - Ha (median) | 75.94 (12.96) | 147.37 (7.37) | 0.007 |
| Number of houses (median) | 1726 (240) | 4384 (219.2) | 0.139 |

\*p-value: Wilcoxon test

A single citizen scientist found an average of 70.68% free-roaming dogs compared to the total dogs detected by the proficient investigator in a community. That proportion increased to 88.7% for two citizen scientists and 95.67% for three citizen scientists. Finally, four citizen scientists were able to detect 100% of the free-roaming dogs, so we established this as the optimal number of searchers to accurately determine the number of free-roaming dogs using GSV (Table 2). In instances, when four citizen scientists in the same community detected more of the free-roaming compared to the findings of the proficient investigator, that particular count was designated as the gold standard for the respective community.

**Table 2.**
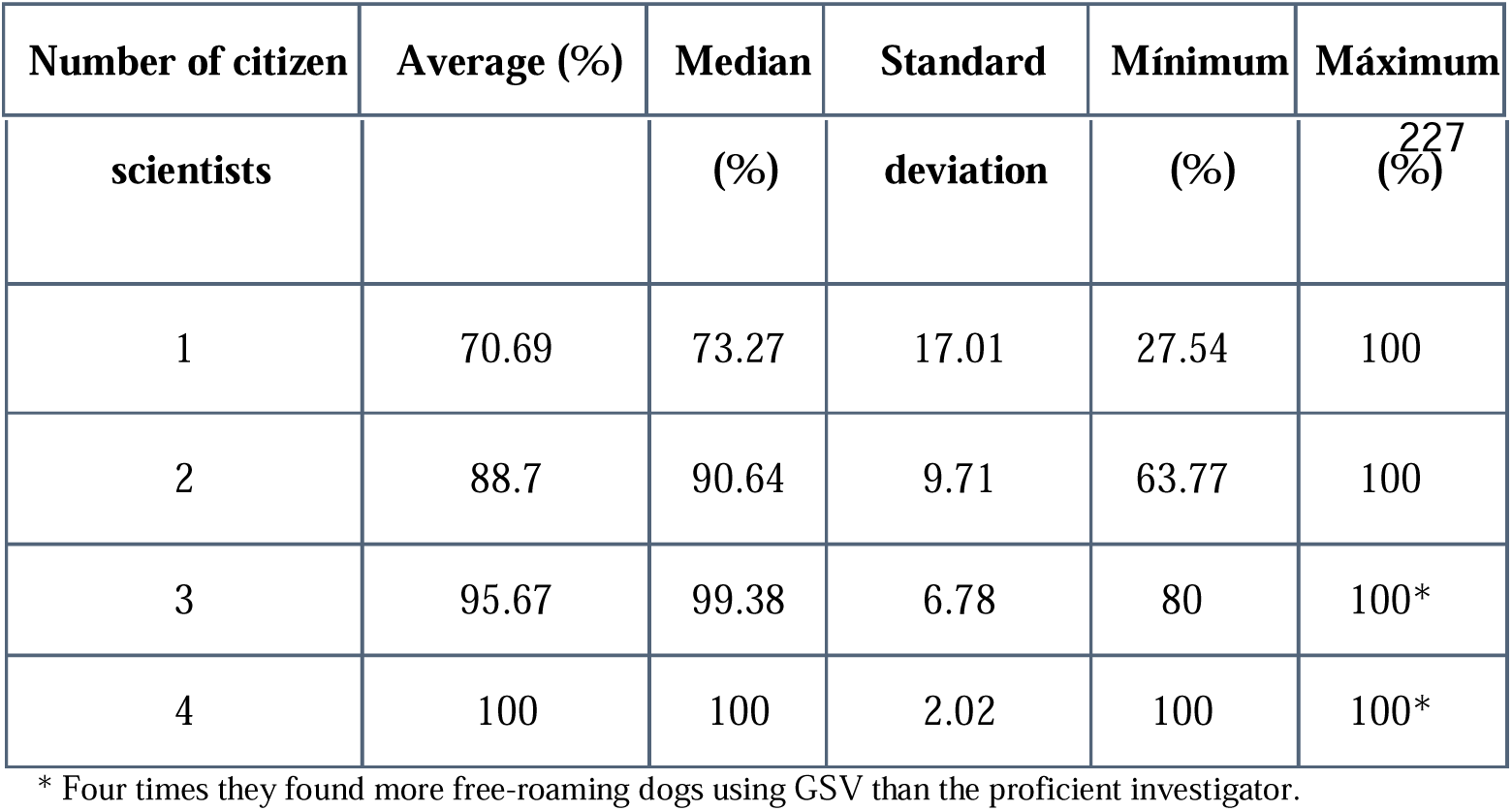
Detectability of free-roaming dogs using GSV relative to the number of searchers/citizen scientists.

| Number of citizen | Average (%) | Median | Standard | Mínimum | Máximum |
| --- | --- | --- | --- | --- | --- |

| scientists |  | (%) | deviation | (%) | <sup>227</sup><br>(%) |
| --- | --- | --- | --- | --- | --- |
| 1 | 70.69 | 73.27 | 17.01 | 27.54 | 100 |
| 2 | 88.7 | 90.64 | 9.71 | 63.77 | 100 |
| 3 | 95.67 | 99.38 | 6.78 | 80 | 100* |
| 4 | 100 | 100 | 2.02 | 100 | 100* |

\* Four times they found more free-roaming dogs using GSV than the proficient investigator.

We found strong and very strong correlations between the response variable and all the predictor variables (Table 3). Free-roaming dogs detected by GSV and owned dogs with free access to the street reported in the surveys presented a strong correlation with a coefficient of 0.73 (p<0.001).

**Table 3.** Pearson correlation coefficients between the dog populations reported in the 2016 surveys and the dog population estimated by GSV during the study in the communities.

| Pearson Correlation |  |  |  |  |  |  |  |
| --- | --- | --- | --- | --- | --- | --- | --- |
| Variables | Dogs found in GSV | Free-roaming dogs surveys | Free-roaming dogs or that walk without a leash with their owners surveys | Total population of dogs survey | Kilometers of travel | Area (Ha) of the community | Number of houses in the community |
| Dogs found in GSV | 1 | 0.73* | 0.78* | 0.75* | 0.91* | 0.74* | 0.83* |
| Free-roaming dogs surveys | 0.73* | 1 | 0.92* | 0.79* | 0.83* | 0.80* | 0.80* |
| Free-roaming dogs or that walk without a leash with their owners surveys | 0.78* | 0.92* | 1 | 0.94* | 0.91* | 0.76* | 0.80* |
| Total population of dogs surveys | 0.75* | 0.79* | 0.94* | 1 | 0.90* | 0.69* | 0.80* |
| Kilometers of travel | 0.91* | 0.83* | 0.91* | 0.90* | 1 | 0.85* | 0.92* |
| Area (Ha) of the community | 0.74* | 0.80* | 0.76* | 0.69* | 0.85* | 1 | 0.86* |
| Number of houses in the community | 0.83* | 0.80* | 0.80* | 0.80* | 0.92* | 0.86* | 1 |

\* Correlation coefficient with a p. value <0.01

### Predicting the population of free-roaming dogs

To select the variables included in the model, we assessed multicollinearity by analyzing the degree of correlation between the potential predictor variables and the variance inflation factor (VIF). A VIF value higher than 10 was used to determine multicollinearity among the regression variables. The following correlations were positive and significant: kilometers and dogs detected by GSV (r=0.91; p<0.001); kilometers and area (r=0.85; p<0.001); and kilometers and the number of houses (r=0.95; p<0.001). In urban communities, there was a clear linear association between the number of free-roaming dogs detected with GSV and the number of free-roaming dogs reported in household surveys. However, this association was completely lost in periurban areas (Fig. 4) A negative binomial regression was used to build the model, and the number of dogs detected by GSV (discrete) and type of community (categorical: urban/periurban) were used for modeling. For model selection, we used Akaikes information criteria (AIC) and the significance of the variables included in our model. As shown in Table 4A, in urban areas, the univariate negative binomial models showed a significant statistical association between the number of free-roaming dogs reported in household surveys and several variables: the number of dogs detected with GSV, the number of houses in the community, the total lengths of roads, and the area of the community—all with a significance level of p<0.001. In the final multivariable model, which included only dogs detected with GSV and the number of houses, only the variable for dogs detected with GSV retained statistical significance (p=0.046). In periurban areas, the data tell a very different story. As shown in Table 4B, the univariate negative binomial models point out that only the number of houses in the community had a significant statistical association with the number of free-roaming dogs reported in household surveys (p=0.029). Interestingly, in the final multivariable model, which included both dogs detected with GSV and the number of houses, only the variable for the number of houses retained its statistical significance (p=0.013).

**Table 4A.** Association coefficients for free-roaming dogs using Google Street View in urban communities of Arequipa, Peru.

| Variable | Univariate model,<br>rate ratio (95% CI) | p-value | Multivariable model,<br>rate ratio (95% CI) | p-value |
| --- | --- | --- | --- | --- |
| Dogs found using GSV | 1.03 (1.02 - 1.04) | <0.001 | 1.03 (1.00 - 1.07) | 0.046 |
| Number of houses | 1.00 (1.00 - 1.01) | <0.001 | 1.00 (1.00 - 1.00) | 0.751 |
| Distance traveled (Km) | 1.39 (1.20 - 1.65) | <0.001 | – | – |
| Area (Ha) | 1.08 (1.04 - 1.14) | <0.001 | – | – |

**Table 4B.** Association coefficients for free-roaming dogs using Google Street View in peri-urban communities of Arequipa, Peru.

| Variable | Univariate model,<br>rate ratio (95% CI) | p-value | Multivariable model,<br>rate ratio (95% CI) | p-value |
| --- | --- | --- | --- | --- |
| Dogs found using GSV | 0.99 (0.98 - 1.01) | 0.681 | 1.00 (0.99 - 1.01) | 0.252 |
| Number of houses | 1.00 (1.00 - 1.00) | 0.029 | 1.00 (1.00 - 1.00) | 0.013 |
| Distance traveled (Km) | 1.05 (0.84 - 1.32) | 0.660 | – | – |
| Area (Ha) | 0.98 (0.94 - 1.03) | 0.475 | – | – |

The estimated equations to calculate the predicted value of dogs in the surveys using the variables Dogs found using GSV and a Type of Zone (urban versus periurban) are the following:

Y = Dogs with free access to the street reported in surveys Y ∼ Negative Binomial (λ)

log(λ) = β0 + β1 *\* (#* Dogs found in GSV) + β2 * (Type of Zone Urban) + β3 *\* (#* Dogs found in GSV)*(Type of Zone Urban)

## Discussion

The results of this study in 26 urban and periurban communities of Arequipa, Peru, showed a strong correlation between the population of dogs with free access to the street reported in door-to-door surveys versus free-roaming dogs detected using GSV. The prediction model with the best fit included the number of dogs detected with GSV, the number of households, and the interaction with the type of community (urban or periurban), highlighting the heterogeneous ecology of free-roaming dogs within cities and evidencing that this new method is viable, at least in well-urbanized areas (35,36). For clarity, we presented separate multivariable models for urban and periurban communities. In periurban communities, predictability was almost null, maybe because the sample size in peri-urban areas was relatively small. Additionally, the virtual exploration of periurban communities using GSV images was less comprehensive relative to urban areas due to the unavailability of GSV images in certain areas or streets. This observed sparsity may be caused by the challenging periurban terrain that impedes the Google vehicles from traveling some streets to capture images, an issue that has been reported by others (37). However, it is also possible that the ecology of free-roaming dogs is very different in peri-urban areas. Urban and periurban areas differ in many aspects that impact the epidemiology of rabies and other infectious diseases, from terrain quality, to access to health services and other public goods, to socioeconomic status and animal and human demography (38–40). Differences in all these factors could impact the ecology of free-roaming dogs explaining the contrast observed in this study.

The number of citizen scientists searching dogs by GSV proved to be a crucial component. We found that four trained citizen scientists accurately counted free-roaming dogs. Interestingly, the same number of citizen scientists has already been reported as appropriate for searching vulture nests using GSV (26). In the present study, four citizen scientists found almost always 100% of the dogs detected by the most proficient investigator using GSV and this percentage decreased with fewer citizen scientists. The low detection of free-roaming dogs when fewer citizen scientists performed the search in the same community could be due to different causes, such as variability in performing the virtual tour, low visual acuity, differences in attention span, or lack of motivation to carry out a full search, this has been observed in other studies (41,42).

We discovered a positive and significant correlation between the number of dogs detected with GSV, the number of dogs with unrestricted street access, and the number of dogs that walk with their owners without a leash, both of which were reported in household surveys. These populations of free-roaming dogs are the most important with regards to zoonoses; they have the highest contact rates between dogs and usually the highest pathogen transmission rates. Also they are less likely to receive preventative services (e.g., vaccination, deworming) because they are less available or are more difficult to handle. Therefore, free-roaming dogs should be the most important targets of zoonotic disease control program. For instance, the population of free-roaming owned dogs impacts the duration of rabies outbreaks (43). When comparing the number of dogs detected with GSV with those reported in door-to-door surveys, the results varied among communities. At the community level, counts with GSV overestimated and underestimated the free-roaming dog populations; however, the overall estimate of the study area was very accurate. Other authors have noted difficulties when comparing alternative methods to surveys; for example, counting dogs with drones underestimates the total dog population because dogs may hide under trees or cars, avoiding detection by the drones (41). On the contrary, we observed that GSV, generally, allows detection of dogs under trees and even cars. A potential problem with observing and registering dogs is double counting. In Tulum, Mexico, using the modified method of photographic capture-recapture, they reported that one dog double counted reported (0.32%) (45). In our study, 1.72% of total dogs detected were double counted. The authors of the Tulum study noted that their method could lead to underestimation of the free-roaming dog population, since they had to sample a fraction of all streets to conduct their transects (45).

Other methods involving direct observation have been evaluated. In Ecuador, Cardenas et al. compared two methods of estimation of free-roaming dogs on transects (capture-recapture and distance sampling) (15). They trained volunteers, obtaining twice as many dogs using the distance sampling method with respect to the capture-recapture method in urban areas. They also found that low light at the time of surveys affected the accuracy of the individual identification and errors in calculating the distance between the animal and the observer caused underestimation of the density of dogs/km² in rural areas. Compared to those methods, counting dogs with GSV requires less time and logistical material. We found the GSV performance was better in urban areas compared to periurban areas. Similarly, Cardenas et al. proposed alternative methods for different areas or even combining different methods to increase accuracy (15). The Ministry of Health in Peru has conducted several workshops to train local staff to estimate their dog populations (14), however, the lack of human resources has impeded these activities. As in many other areas in Latin America, in Arequipa, Peru, the dog population has been estimated using the human-to-dog ratio. In 2015, the ratio they used was 10 to 1, while in 2016 it was 5 to 1, and these ratios were applied to all districts in the city, without considering the spatial variability between them. This level of uncertainty and the lack of spatial precision have greatly affected the assessed effectiveness of mass dog vaccination campaigns aimed at eliminating dog-mediated human rabies (33). Using the GSV imagery for estimation of the free-roaming dog population can reduce this uncertainty and even feed into models to determine hot spots of transmission for different diseases transmitted by dogs such as rabies (46).

The method employed in our study showed some limitations. Firstly, the variability of images in GSV due to periodic updates may affect the consistency of dog observations made by citizen scientists. Additionally, the availability of the Street View function in certain areas (e.g., alleys, unpaved secondary roads) may limit the search for free-roaming dogs. Furthermore, uncertainties regarding the timing and frequency of image capture introduce unknown intervals for analysis. It is important to acknowledge that our count of free-roaming dogs using GSV includes unowned dogs, unlike surveys conducted in 2016 that only registered dogs reported by owners (47). It is important to note that, according to the Ministry of Health of Peru, 90% of free-roaming dogs in cities are owned (48), so our results might not be strongly affected by this unaccounted dog subpopulation However, the stray dog should also be estimated to provide a full picture of the target population for zoonotic disease control programs. Despite these limitations, utilizing GSV presents advantages over traditional methodologies, particularly during circumstances like the COVID-19 pandemic or in resource-limited settings, as it eliminates the need for field trips by trained personnel (12,15,49,50). Moreover, the periodic updating of the Google Street View database (18) enhances the potential for timely updates on free-roaming dog populations across different regions.

The present study demonstrates Google Street View can be a useful tool for obtaining accurate and reliable estimates of the free-roaming dog population. The method also offers several advantages over household surveys, transect counts, or photographic capture-recapture. GSV is already used for construction projects, geological evaluations, ecological studies, and even assessment of chronic diseases (22,25,26,51). Pairing GSV with newer technologies such as machine learning or artificial intelligence could enhance its potential and make it faster and more available for programs without human resources to conduct virtual searches. In addition to obtaining the dog population size, with GSV it is also possible to obtain the exact geographic coordinates of dogs, which facilitates the analysis of their distribution relative to biological events and geographical features. Information from such analysis could feed control programs for various dog-mediated diseases of public health (7,52) as well as diseases threatening other animal species (53–58). The use of GSV imagery for dog population estimation provides a viable approach to address the limited knowledge about free-roaming dogs in urbanized areas. The method requires minimal resources and has the potential to inform effective public health policies and programs addressing free-roaming dogs.

## Acknowledgments

We thank the citizen scientists who voluntarily contributed to the study. We acknowledge the work of the members of the Zoonotic Disease Research Laboratory, One Health Unit at Universidad Peruana Cayetano Heredia for their valuable contributions in collecting a part of the data utilized in this study. Lizzie Ortiz and Raquel Gonçalves provided support by reading this article.

